# Effort cost of reaching prompts vigor reduction in older adults

**DOI:** 10.1101/2023.08.28.555022

**Authors:** Erik M. Summerside, Robert J. Courter, Reza Shadmehr, Alaa A. Ahmed

## Abstract

As people age, they move slower. Is age-related reduction in vigor a reflection of a reduced valuation of reward by the brain, or a consequence of increased effort costs by the muscles? Here, we quantified cost of movements objectively via the metabolic energy that young and old participants consumed during reaching and found that in order reach at a given speed, older adults expended more energy than the young. We next quantified how reward modulated movements in the same populations and found that like the young, older adults responded to increased reward by initiating their movements earlier. Yet, their movements were less sensitive to increased reward, resulting in little or no modulation of reach speed. Lastly, we quantified the effect of increased effort on how reward modulated movements in young adults. Like the effects of aging, when faced with increased effort the young adults responded to reward primarily by reacting faster, with little change in movement speed. Therefore, reaching required greater energetic expenditure in the elderly, suggesting that the slower movements and reactions exhibited in aging are partly driven by an adaptive response to an elevation in the energetic landscape of effort. That is, moving slower appears to be a rational economic consequence of aging.

**Significance statement:** Healthy aging coincides with a reduction in speed, or vigor, of walking, reaching, and eye movements. Here we focused on disentangling two opposing sources of aging-related movement slowing: reduced reward sensitivity due to loss of dopaminergic tone, or increased energy expenditure movements related to mitochondrial or muscular inefficiencies. Through a series of three experiments and construction of a computational model, here we demonstrate that transient changes in reaction time and movement speed together offer a quantifiable metric to differentiate between reward- and effort-based alterations in movement vigor. Further, we suggest that objective increases in the metabolic cost of moving, not reductions in reward valuation, are driving much of the movement slowing occurring alongside healthy aging.

## Introduction

Among healthy people there are stable inter-subject differences in movement speed: some individuals consistently move faster than others [1]. Indeed, a prominent factor that influences movement speed is aging. Older adults walk [2,3], reach [4–6], and make saccadic eye movements [7,8] at a slower speed than younger adults. Why do older people move slower?

In principle, a reduction in vigor may be due to changes in the reward system of the brain [9]. The brain’s ability to accurately predict the value of an upcoming reward depends in part on the integrity of the dopaminergic neurons [10]. Moving faster requires expenditure of greater effort [11–13],and individuals with dopamine deficits exhibit a diminished willingness to exert effort to acquire reward [14,15]. Because the integrity of the dopaminergic system declines with healthy aging [16,17], a reduced sensitivity to reward may be one explanation for why older adults make slower movements [18].

An alternative and perhaps complementary explanation is that as one grows older, it becomes more effortful to make movements. For example, older adults require a greater rate of energetic expenditure to walk at a given speed [2,19,20] or to produce a given power on a cycle ergometer [21,22]. This elevated energetic expenditure is associated with a decrease in efficiency of the mitochondria and contractile elements of muscles [21], as well as an increase in the levels of co-activation of antagonist muscles [20,23,24], both of which occur in aging. Therefore, in addition to a decline in the reward system of the brain, aging coincides with an increase in the energetic requirements of movement. This raises the hypothesis that with aging, moving slower is a rational economic decision in response to a reduced expectation of reward and an increased expectation of effort.

Here, we attempted to quantify this economic landscape of movements in aging. We focused on a reaching task and measured the energy that the elderly expended as they moved their hand from one point to another. We found that the metabolic cost of reaching was indeed greater in the elderly than in the young. That is, the cost of performing a reach at a given speed was objectively larger. Next, we quantified response to reward and found that like younger adults, the older participants responded to increased reward by reducing their reaction time. However, unlike the young, they were less willing to increase their movement speed. Given the increased cost of movement in the elderly, is the reduction in reaction time but not movement duration a rational response to increased reward?

To answer this question, we considered a normative model of behavior from foraging theory in which the objective of actions is to maximize a capture rate: reward minus effort, divided by time. The mathematics confirmed that given their increased effort cost, when faced with the opportunity for greater reward the elderly should respond not by increasing movement speed, but by primarily reducing their reaction times. In contrast, the young should do the opposite, primarily increasing their movement speed rather than altering reaction times.

To test these predictions, we performed a third experiment in which we tried to make the young experience reaching as if they were old: we increased their reach effort costs (with respect to a baseline) and then measured how they responded to increased reward. When their effort costs were elevated, the young responded like the elderly in that they reacted to increased reward by reducing their reaction times, not by moving faster.

Together, our results demonstrate that the cost of reaching is objectively higher in older adults, and that their slower movements may be primarily due to higher effort costs, not impaired reward valuation.

## Results

We sought to understand why healthy aging accompanies a reduction in movement speed. To answer this question, we measured the metabolic cost of reaching in groups of young (18-35 years) and older (66-87 years) adults. In our first experiment, we measured the rate of energy expenditure as participants reached at various speeds. In our second and third experiments, we measured motor responses to varying amounts of reward when reaching with high and low effort costs.

### Cost of reaching is greater in the older adults

We measured the rate of energy expenditure at rest and during reaching. In the baseline period, younger (n = 12; 25 ± 2 years; 6 Females) and older (n = 12; 75 ± 8 years; 6 Females) subjects held the handle of a robotic arm and remained still (Fig. 1A). The resulting rate of energy expenditure *ė*_*o*_ was not significantly different between the two groups (t-test, young = 77 ± 15W, old = 69 ± 16W, p = 0.241). Thus, at baseline, energetic rates were comparable between our sample of young and older people.

**Figure 1.**
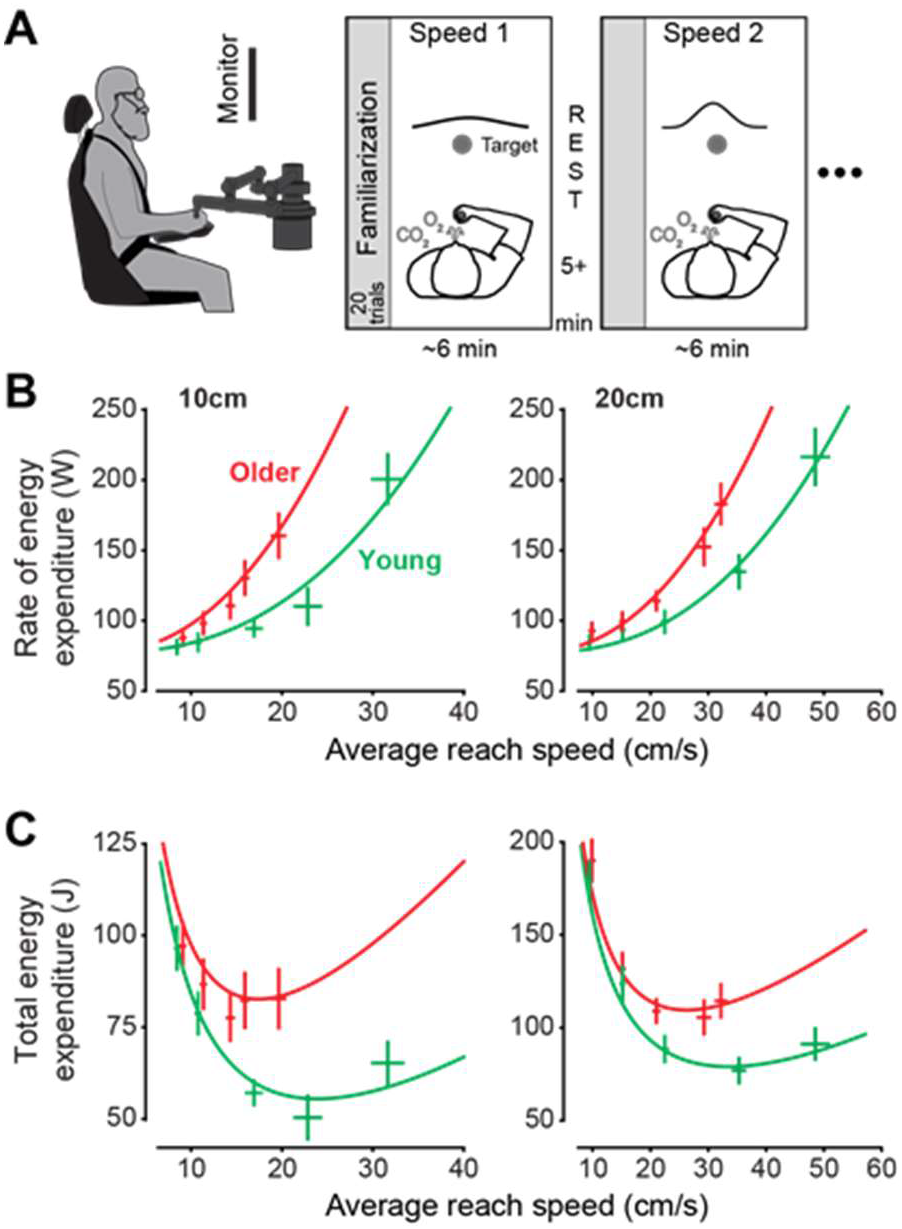
Experiment 1: Reaching is energetically more costly in older adults. A) Participants controlled a cursor presented on a monitor by moving a robotic manipulandum with their right hand in the horizontal plane at various prescribed speeds and distances. B) Rate of energy expenditure increased with reach speed at a given distance, but this cost was greater in the older adult group. Curves represent best fit from Eq. 1: *a*_*young*_=77.33 [67.84 85.49]W, *a*_*older*_=77.52 [60.14 92.38]W; and *b*_*young*_= 114.67 [44.60 226.23], *b*_*older*_= 151.44 [49.01 334.30], *i*_*young*_=1.23 [0.83 1.67] (mean [95%CI]), *i*_*older*_=0.88 [0.52 1.40]; *j*_*young*_=2.44 [1.85 3.15], *j*_*older*_=2.17 [1.39 3.55]. C) Total energy expenditure (cost of reaching). Vertical and horizontal error bars represent ±SEM.

As subjects reached, the rate of energy expenditure *ė*_*r*_ increased with average reach speed (Fig. 1B, β = 0.5373, 95%CI [0.48, 0.59], p < 0.001). However, the older adults required a greater rate of energy expenditure to reach at a given speed (main effect of age, β = 0.1813W, 95%CI [0.04, 0.31], p = 0.012). For example, the metabolic cost of reaching a 10 cm distance at an average speed of 20 cm/s was 35% greater for the older adults than the young (57 Joules in the young vs. 77 Joules in the elderly).

To functionally represent the relationship between energy expenditure, target distance *d*, and reach duration *T*_*r*_, we used a previously described formulation [13]:

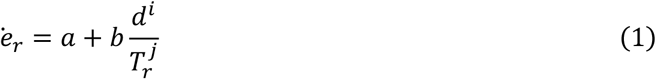

This equation captured the measured data well (Fig. 1B, r_young_ = 0.85 [0.78 0.90], r_old_ = 0.67 [0.59 0.75]). By integrating Eq. 1 as a function of time, we computed the total energy that was expended to reach a distance *d*. The result was a concave-upward function of reach duration *T*_*r*_ (Fig. 1C). This energy cost and was consistently larger in the older adults than the young:

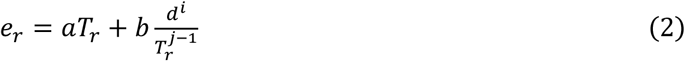

What might be the reason for this increased cost in the elderly? The simplest possibility is that perhaps the older adults were burdened by a heavier arm. To investigate this, we estimated the mass of the arm in each participant as a function of their total body mass, age, and sex using previously published observations [25,26]. We found no significant differences in estimated mass between younger and older adults for the upper arm (m_young_ = 1.75 ± 0.33kg, m_old_ = 1.77 ± 0.39, p = 0.898), lower arm (m_young_ = 1.00 ± 0.23kg, m_old_ = 1.16 ± 0.35, p = 0.197), or hand (m_young_ = 0.39 ± 0.08kg, m_old_ = 0.43 ± 0.12, p = 0.331). Thus, we could not attribute the elevated energetic cost of reaching in older adults to a heavier limb.

Another possibility is that in the older adults, movements were jerkier, requiring costly corrections [27]. Indeed, movements of the older adults were less accurate than the young, as illustrated by endpoint errors along the axes normal and tangential to the path to the target (Fig. S1A-D, normal: β = 0.330mm, 95%CI [-0.14, 0.81], p = 0.168, tangential: β = 3.503mm, 95%CI [1.33, 5.68], p = 0.003). Perhaps these errors led to more corrective movements when approaching the target, subsequently resulting in greater effort costs. A way to probe the extent of corrective movements is to calculate the smoothness of the movement as the sum of squared jerk during the reach. We found that while the sum of squared jerk increased with faster movements (β = 1345012.34 (m/s^3^)^2^, 95%CI [1143804.86, 1546045.69], p < 0.001), there was no effect of age (β = -34805.62, 95%CI [-82692.01, 13245.85], p = 0.152) (Fig. S1E,F).

In summary, reaching consumed a greater amount of energy in the elderly, and tended to be less accurate.

### Like the young, older adults respond to reward by reducing reaction time, but they are less willing to increase their reach speed

In Exp. 2 we tested how the old and the young responded to reward. Young (n = 20; 26 ± 4 years; 10 Females) and older (n = 20; 72 ± 6 years; 10 Females) participants moved a cursor in an out-and-back motion toward a very large target, indicated by a 100° arc (Fig. 2A)[28]. The study design included the very large targets because we wanted a task in which reach endpoint accuracy did not play a significant role.

**Figure 2.**
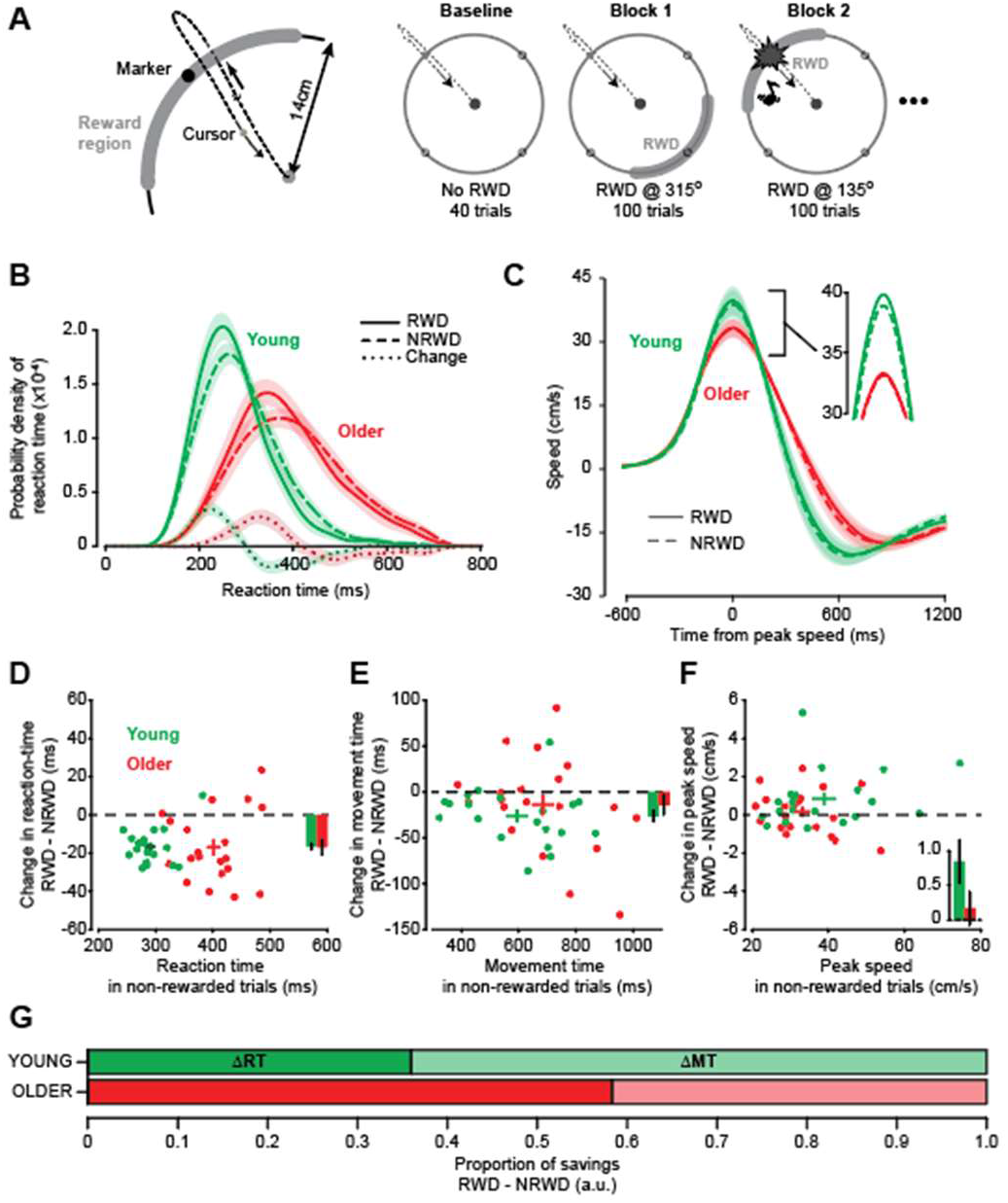
Experiment 2: Reward quickens reaction time in young and old. A) Participants performed out-and-back reaches to alternating targets projected along a ring 14cm from the home circle. The desired quadrant was cued with a marker centered at the middle of the quadrant. Visual feedback of the cursor was removed during the outward portion of the movement and was re-displayed during the return portion of the reach once the hand was at 9cm from the home target. The protocol consisted of a baseline period with no reward followed by four experimental blocks. Each experimental block had one quadrant paired with a reward (RWD). Audiovisual reward stimulus was delivered upon crossing any region of the 100° target arc. The gray areas indicating reward were not visible to the participant, but are presented in the figure to convey which quadrant was paired with reward. B) We used a non-parametric kernel density estimation method to calculate the probability distribution for each individual when making movements to rewarded (RWD, solid curves) and non-rewarded (NRWD, dashed curves) quadrants as well as a difference (dotted curve) in these distributions at each bin (bin size=5ms). Younger adults (green curves) initiated movements earlier than older adults (red curves), but both groups responded to reward by reacting sooner. C) Effects of reward on movement execution in young (green) and older (red) adults. Young adults made movements toward quadrants paired with reward (RWD, solid curves) with greater peak speed when compared to that same quadrant when not rewarded (NRWD, dashed curves). Older adults reached with a peak speed that was independent of reward status. Inset graph depicts enlarged region highlighted peak speed. D-F) Scatter plot representing the relationship between rewarded (RWD, vertical axis) and non-rewarded (NRWD, horizontal axis) movements according to mean reaction time (D), movement duration (E), and peak speed (F). Dots represent individual participants. Crosses represent the mean for each age group and the length of the bars represent ±SEM. The mean effect of reward for each age group is indicated with the inset bar graph, reported as mean ± SEM. G) Proportion of time savings due to reaction time (RT) and movement time (MT) in young vs. older adults. The proportion of time saved by reacting faster is larger in older adults.

Subjects were free to select their reach speed. Within a given block, one of the quadrants was consistently paired with reward while the remaining three quadrants were always unrewarded. The reward consisted of a short, pleasant tone, a visual flashing of the outer ring, and four points. We had found previously that this feedback was a reasonable proxy for reward as it led participants to invigorate their reaching movements [28]. The only requirement for reward was that the cursor crossed anywhere along the 100° arc of the indicated quadrant.

Older adults took longer to initiate their movements. Their average reaction time was 401 ± 12ms, significantly longer than the reaction times (292 ± 7ms) observed in the young (Fig. 2B, mixed effects model, β = 109.6ms, 95%CI [81.5, 137.6], p < 0.001). In trials toward the rewarded quadrant, both young and older adults responded by reducing their reaction time (Fig. 2D, change in mean reaction time, Δ_young_ = -16 ± 2ms, Δ_old_ = -17 ± 4ms; main effect of reward: β = -16.57ms, 95%CI [-21.1, - 11.9], p < 0.001; reward by age interaction: p = 0.863).

However, in contrast to the consistent and robust effects of reward on reaction times of both older and young adults, we found a reduced effect of reward on reach speed in the older population as compared to young (main effect of reward: β = 0.84cm/s, 95%CI [4.88, 12.0], p = 3.43e-6; reward by age interaction: β = -0.68cm/s, 95%CI [-11.8, -1.8], p = 0.0081). Whereas young adults reached faster toward the rewarding target (Fig. 2C & F, Δ_young_ = 0.84 ± 0.182cm/s), reward did not significantly increase reach speed in the older adults (Δ_old_ = 0.16 ± 0.181cm/s). Similarly, while reward reduced movement durations in both groups (β = -26.06ms, 95%CI [-39.1, -13.0], p < 0.001), older adults reduced their movement duration to a lesser extent than young adults (Fig. 2E, Δ_young_ = -26.06 ± 6.649ms, Δ_old_ = -13.89 ± 6.619ms; reward by age interaction: β = 12.16ms, 95%CI [-6.22, 30.6], p = 0.1948).

Both young and older adults altered their movements to save time and acquire reward faster. Yet each group used a different strategy. The young relied primarily on faster movements, which accounted for approximately 65% of the total time savings on average. In contrast, older people relied primarily on faster reaction times, which drove nearly 60% of the time savings (Fig. 2G).

In summary, both the young and old reduced their reaction time when they were reaching toward a rewarding target. However, older adults were less willing to increase the speed of their movements in response to reward.

### A greater effort cost should lead to slower movements and longer reaction-times

To better understand these results, we tried to ask how the elderly should alter their reaching movements in response to increased reward. Is their reluctance to increase speed but their willingness to reduce reaction time a rational response to age-related changes in effort costs?

If we assume that the reward promised at the end of the movement interacts with the effort that must be expended to make that movement, the result is a utility that can, in principle, specify the optimal movement. A normative form of this utility [29] is employed in the field of optimal foraging: utility (J) is defined as the capture rate, i.e. the difference between the reward attained and effort expended, divided by time required to obtain that reward [30–32]. Earlier work has shown that this formulation makes testable predictions regarding how changes in the reward and effort landscape should affect patterns of movement [33]. Here, we used this idea to ask the following question: given that the older adults are burdened by a greater energetic cost of reaching, and suffer from greater inaccuracy, how should they reach?

When a potentially rewarding target is presented to a subject, the duration of time that passes before acquisition of reward includes both the reaction time *T*_*o*_, and the duration of the movement *T*_*r*_. Let reward magnitude be specified by α, and use the capture rate to define the utility of the reach:

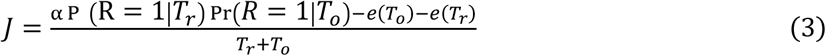

In the above expression, Pr*(R* = 1|*T*_*r*_) is the probability that a movement of a given duration would succeed in acquiring reward, *e*_*r*_ is the metabolic cost of reaching (Eq. 2), and *e(T*_*o*_) is the metabolic cost of waiting before starting the movement (i.e., the cost incurred by the reaction time). To approximate this cost, we assumed *e(T*_*o*_) included the metabolic cost associated with holding still, which we measured via baseline metabolic rates, *ė*_*o*_:

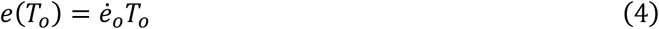

The term Pr*(R =* 1|*T*_*r*_) captures the speed-accuracy trade-off, i.e., the probability of acquiring the reward given that the movement was of duration *T*_*r*_. To estimate Pr*(R =* 1|*T*_*r*_), we used the endpoint accuracy data that we had measured in our subjects for a 10 cm movement across the range of speeds and fit a logistic function with parameters *b*_0_ and *b*_l_ in Eq. 5. As expected, for a given movement duration, older adults were less accurate than the younger adults (Fig. 3B).

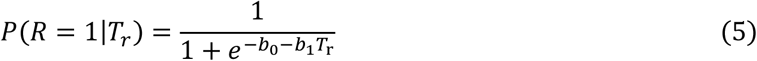

**Figure 3.**
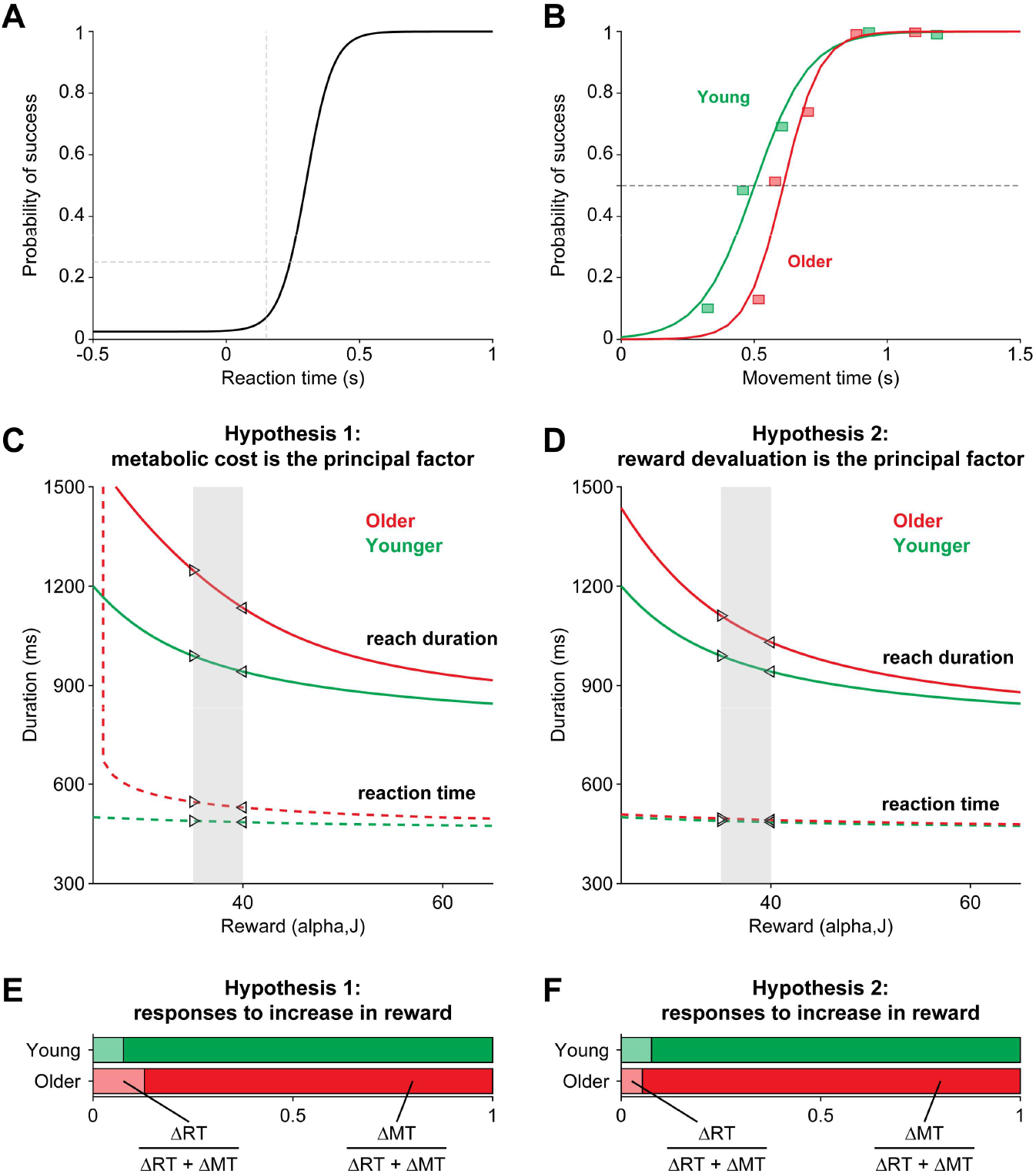
Model: Effort costs slow movement and reaction time. A) Logistic function representing the speed-accuracy tradeoff for reaction times. B) Logistic curves fitted to endpoint data of older (red) and younger (green) adults representing the speed-accuracy tradeoff for movement times. For a given reach duration, older adults saw a lower probability of success than the young. C) Effect of age on optimal reaction times (dotted lines) and movement times (solid lines) across arbitrary reward values, using (Eq. 3). Fitted metabolic parameters (Fig. 1B) and associated speed-accuracy curves were used for the young and older curves. D) Effect of reward valuation on optimal reaction times and movement times. Green curves represent optimal solutions based on the younger adult metabolic fits, accuracy, and nominal reward valuation (k=1). Red curves are optimal solutions with the same effort costs, but older adult accuracy and reduced reward valuation (k=0.8) E & F) The proportions of time saved (Δ*RT/(*Δ*RT +* Δ*MT)*) due to reducing reaction time or movement time for an arbitrarily selected increase in reward from 45J to 50J (gray region in C & D). Compared to young, older adults should allocate a larger proportion of time-savings to reducing reaction times due to higher metabolic costs (E). If older adults were instead valuing reward less, their proportion of time-savings from reaction time should instead be lower (F).

We used the term Pr*(R =* 1|*T*_*o*_) to represent the fact that reaction times affect accuracy movement direction, with the probability of success increasing with longer reaction times [34,35]:

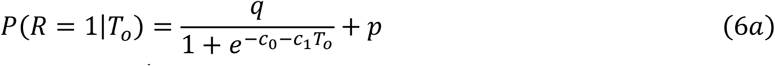

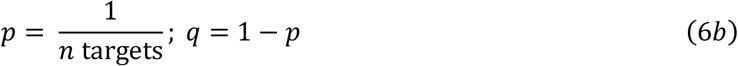

We considered a scenario where there are *n* = 40 potential targets (i.e., the maximum number of non-overlapping 0.8 cm radius targets that can fit around a 10 cm radius circle), and thus an early reaction time would result in a chance probability (*p*) of a reach towards the correct target (Eq. 6b; Fig. 3A). Next, we computed the reaction times 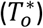 and movement durations 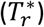 that maximize this utility:

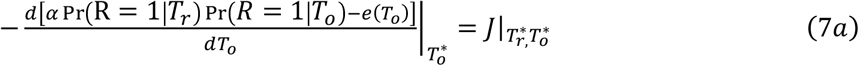

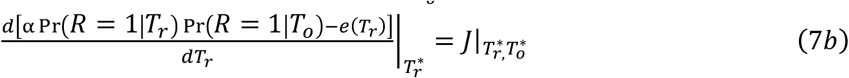

These equations made sensible predictions: in the face of an increased metabolic cost of reaching, and increased inaccuracy, the best policy is to reduce reach speed, and increase reaction time (Fig. 3C). Importantly, these results were maintained even when using equivalent speed-accuracy tradeoffs between the young and older adults, as well as across a range parameter values (Eq. 5). Thus, an increased cost of reaching alone is sufficient to slow movements.

Next, we used the equations to examine the results of Exp. 2. Under hypothesis 1, the elderly move slower than the young primarily because of their increased effort costs. Under hypothesis 2, elderly move slower than the young primarily because they value reward less. Can the results of Exp. 2 help us dissociate between the predictions of these two alternatives?

In the elderly, energetic costs were greater than the young, and movements suffered from greater inaccuracy. However, the energetic cost of reaction time was the same (because in our two groups, the baseline metabolic costs were not different). To compute the predictions of hypothesis 1, we inserted the measured metabolic costs and accuracy into Eqs. (7) and computed how reaction time and movement duration should change in response to a given change in reward via the following ratios: 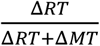 and 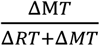.

We found that according to hypothesis 1 (increased effort cost), in response to increased reward both the elderly and the young should reduce their movement duration and reaction times, but because moving faster costs more for the elderly, the elderly should focus more of the change in their reaction times (Fig. 3E). That is, if the increased effort costs are the main issue for the elderly, then they should respond to reward by reducing their reaction time, not movement time.

In hypothesis 2 (reduced sensitivity to reward), we imagined the counterfactual condition that the old and the young had the same effort costs of reaching. Thus, under this hypothesis the elderly moved slower because they valued the reward less. In this case, the two groups differed not in terms of their effort costs, but because of differences evaluation of reward and differences in accuracy (Fig. 3D). We imagined that in the elderly, reward *α* was devalued (represented by *kα*, where *k* < 1), reach accuracy was as measured, but the cost of reaching was the same as in the young. Reward devaluation was sufficient to produce the reduced reach speeds and longer reaction times in the elderly (Fig. 3D). However, hypothesis 2 predicted that in response to increased reward, both the elderly and the young should reduce their movement duration and reaction times, but because increasing movement speed now costs the same in the two groups, the elderly should focus more of the change in their reach speed, not reaction time (Fig. 3F). Thus, according to hypothesis 2, if the slower movements in older adults were primarily a consequence of diminished reward valuation, then compared to the young, the elderly should shorten their reach duration to a greater extent.

In summary, our modeling suggests that the results of Exp. 2 are consistent with Hypothesis 1 and not Hypothesis 2. That is, when effort costs of a reach are increased, it is rational to respond to increased reward by primarily reducing reaction times, not movement time.

### Increasing the effort cost of reaching in the young makes them respond to reward like the elderly

The inference that results from the model is that the elderly may be reaching slower principally because they are burdened with increased effort costs. But to make a causal link between the increased effort costs and their response to reward, we thought of a third experiment: make the young experience the effort costs of the elderly and see if they too would respond similarly to increased reward.

In Exp. 3, we explicitly manipulated the effort cost of reaching in the young. A new group of young participants (n = 20; 23 ± 4 years; 10 Females) completed a protocol like the one detailed in Exp. 2. In Exp. 3, participants completed the paradigm twice – once with low effort (0 kg), and once with high effort (∼3.63kg/8lbs physical mass added to the robotic arm) (Fig. 4A). The order of the effort conditions was randomized and counterbalanced across participants, with 5-10 minutes of rest between each condition. Across the low and high effort conditions, each participant experienced a total of 160 baseline trials, 200 rewarded trials, and 600 nonrewarded trials.

**Figure 4.**
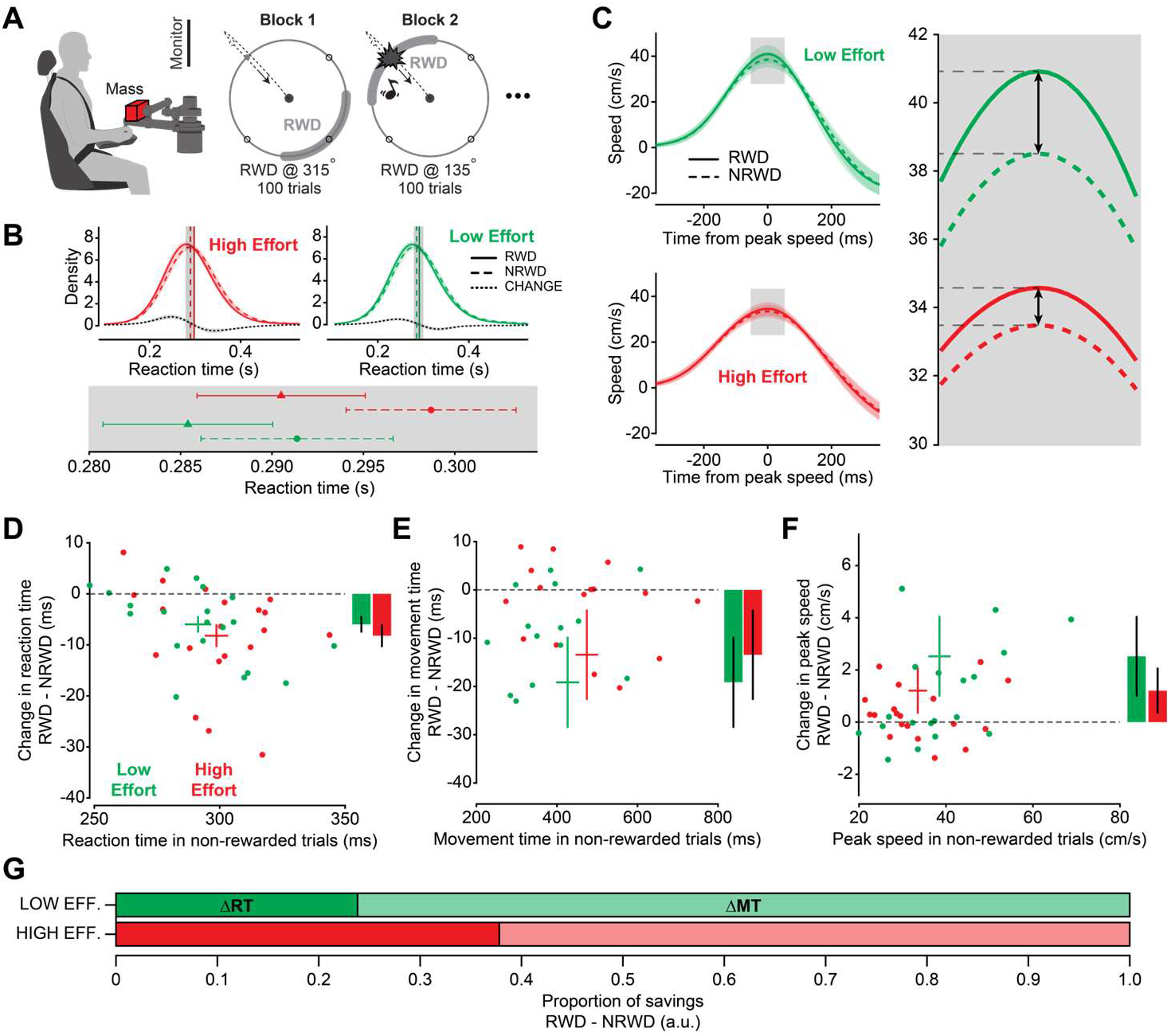
Experiment 3: Effort slows movement and reaction times in younger adults, and mitigates effect of reward on movement speed. A) Design for experiment 3. Participants performed out-and-back reaches to alternating targets projected along a ring 10cm from the home circle. The paradigm was similar to that of experiment 2, except visual feedback of the cursor was maintained for the duration. Participants performed this protocol twice, once with no added mass (0 kg) to the handle of the robot, and once with 3.63kg/8lbs added mass. B) Non-parametric kernel density estimation for the probability distribution reaction times when making movements to rewarded (RWD, solid curves) and non-rewarded (NRWD, dashed curves) quadrants as well as a difference (dotted curve) in these distributions at each bin (bin size=5ms). Low effort (green curves) movements were initiated earlier than high effort (red curves), reward reduced reaction times in both groups. C) Effects of reward on peak speed in low (green) and high (red) effort conditions. Speeds requiring low effort were overall faster than high effort. Rewarded movements had higher peak speeds regardless of effort (RWD, solid curves) compared to non-rewarded (NRWD, dashed curves). D-F) Scatter plot representing the relationship between non-rewarded (NRWD, horizontal axis) and the difference between rewarded and non-rewarded (RWD-NRWD, vertical axis) reaction time (D), movement duration (E), and peak speed (F). Dots represent individual participants. The intersection at each cross represents the mean for each age group and the length of the bars represent ±SEM. The mean effect of reward for each age group is indicated with the inset bar graph, reported as mean ± SEM. G) Proportion of time savings due to reaction time (RT) and movement time (MT) in low vs. high effort (exp. 3). The proportion of time saved by reacting faster is larger when effort is higher.

Reward had a robust main effect on all kinematic parameters, demonstrating a generalized “speeding-up” of movement. Under constant effort conditions, reward reduced reaction time (β = - 5.94ms, 95% CI [-8.71, -3.16], p = 2.72e-05), increased peak velocity (β = 2.50cm/s, 95% CI [2.14, 2.86], p < 2e-16), and reduced movement duration (β = -18.92ms, 95% CI [-24.80, -13.04], p = 5.73e-10) (Figs. 4B-F).

Effort also had a significant main effect on all movement characteristics, tending to slow-down reaches (Figs. 4B-F). For a given reward state, on average, effort increased reaction time (β = 7.44ms, 95% CI [5.48, 9.40], p =1.08e-13), decreased peak velocity (β = -5.00cm/s, 95% CI [-5.26, -4.75], p < 2e-16), and increased movement duration (β = 48.96ms, 95% CI [44.80, 53.12], p < 2e-16) (Figs 4B-F).

When responding to rewarded targets, subjects reduced their reaction time by Δ_low_ = 5.94 ± 1.413ms in the low effort condition, less than the Δ_high_ = 8.22 ± 1.413ms reduction that was present in the high effort condition. In the high effort condition, compared to the low effort conditions, subjects quickened reaction times just as much, if not more to reward (reward by effort interaction: β = -2.28ms, 95% CI [-6.20, 1.64], p = 0.254) (Figs. 4B & 4D).

In contrast, a significant interaction modulated peak velocity’s response to reward in the presence of effort (reward by effort interaction: β = -1.29cm/s, 95% CI [-1.80, -0.78], p = 3.16e-06). This suggests that the increase in reach speed when responding to reward was mitigated by effort. In the low effort condition, peak velocity increased by Δ_low_ = 2.50 ± 0.184cm/s towards a rewarded target as compared to its nonrewarded counterpart (38.51cm/s NRWD compared to 41.01cm/s RWD). In the high effort condition, the increase in peak velocity was smaller, at Δ_high_ =1.21 ± 0.184cm/s, when compared to its nonrewarded counterpart (33.507cm/s NRWD compared to 34.72cm/s RWD) (Figs. 4C & 4F). Similarly, in response to reward, movement duration reduced to a greater extent in the low effort (Δ_low_ =18.92 ± 2.998ms) compared to the high effort condition (Δ_high_ =14.62 ± 2.996ms), although the interaction did not reach significance (Fig. 4E, reward by effort interaction: β = 4.30ms, 95% CI [-4.01, 12.61], p = 0.311). Thus, the increase in movement speed when responding to reward was mitigated by effort.

More effortful movements in younger adults seemingly shift reward sensitivity towards reaction time responsiveness, possibly because reducing reaction time incurs a lower energetic cost than reducing movement duration. These results mirror what was seen with older adults and our theoretical predictions, suggesting that to obtain reward when movement is effortful, increases in movement vigor begin to favor faster reaction times over faster movement times. (Fig. 4G)

## Discussion

We found that the metabolic cost of reaching was elevated in older adults, implying that it was energetically advantageous for them to move slower. But is this objective increase in effort cost a causal factor in the slowing of movements in older adults? To explore this question, we presented young and older subjects an opportunity to acquire reward. In response, both groups decreased their reaction times. However, when reaching towards rewarding stimuli, only the young adults increased their reach speed. In a third experiment, we tried to make the young experience the effort cost of moving like the elderly. We increased the effort cost of reaching in the young by adding a mass to their hand, and again measured their response to reward. When the effort requirements of reaching was increased, the young responded to reward like the older adults: principally through reduced reaction time, not movement time. Taken together, our results suggest that the reduced movement speed in the older adults, as well as their reluctance to modulate this speed in response to increased reward, may primarily reflect their increased effort cost of reaching.

### Metabolic rate of reaching is elevated in older adults

We found that like the young [30], in the older adults energetic expenditure of reaching grows larger with distance and speed. However, for a given speed and distance, older adults expend greater amounts of energy to reach than the young. Similar age-dependent findings have been observed in walking. As we age, we adopt slower walking velocities which are correlated with an overall greater energetic costs [2,19,36].

The elevated cost of reaching in older adults may arise from several factors. The skeletal muscle mitochondria may experience a reduction both in their capacity to generate the needed ATP as well as a reduction in the efficiency with which they convert oxygen to ATP. Coen et al. [37] showed that both mitochondrial capacity and efficiency are reduced with age and correlate with preferred walking speed; reduced mitochondrial function predicted slower preferred walking speeds. Older adults may have also made reaches with greater levels of muscle coactivation, despite extensive familiarization with the task. Increased coactivation may also lead to greater energetic costs [38,39].

Here we considered absolute energetic costs, but it is possible that individuals consider costs relative to their aerobic capacity. Similar to absolute energetic expenditure, aerobic capacity has also been demonstrated to be lower in older adults [37,40]. Thus, irrespective of both the demonstrated age-related increases in absolute energy cost or the possible reduced aerobic capacity, we would predict slower movements and a reduced willingness to respond to reward.

Our results do not exclude the possibility that there is a subjective, age-dependent inflation in the cost of effort, an additional explanation as to why older adults were not willing to adjust their movement speed. While the dopaminergic midbrain has long been a target for the coding of reward value [10,41], there is more recent evidence suggesting that dopamine release rises in anticipation of higher task effort [42]. Wardle et al. [43] were able to identify a positive association between an individual’s level of activity in dopaminergic regions with their willingness to exert effort for a given reward. Similarly, individuals with decreased dopaminergic tone, such as those with Parkinson’s Disease (PD), show a heightened sensitivity to effort [14,15,44]. Behaviorally, dopamine release in the moments before onset of a movement increases speed of the ensuing movement [45], and greater amounts of dopamine are released during movements that require greater effort [42]. In older adults, this dopaminergic midbrain region has been shown to decline in activity as a function of aging [16,46], suggesting that both reward valuation and the willingness to invest effort may be impaired, ultimately leading to reduced movement vigor. Overall, our results implicate effort costs but cannot dismiss contribution from reduced reward valuation.

### Individuals relied on reaction time to obtain the more effortful reward more quickly

When reaching costs increased – either through higher metabolic effort in older adults or through increased mass on the arm in younger adults – we observed a change in the strategy to obtain reward. Though reaction times and movement speeds were on average slower in higher effort conditions [1,47,48], individuals decreased reaction times to a greater extent and mitigated increases in movement speed, and thereby effort, for reward. If we consider the baseline metabolic rate (Eq. 4), then reacting faster saves energy from being wasted through homeostatic processes while still reducing the delay in attaining reward. On the contrary, stunting the increases in movement speed for reward when the effort associated with moving is high will save additional metabolic energy (Figs. 1B & 1C).

These results are partially corroborated in a Parkinson’s study from Kojovic et al. [49] in which they investigated performance in a rewarded simple reaction time task for patients on and off dopaminergic medication. When successful performance was monetarily rewarded, PD patients and healthy controls quickened reaction time irrespective of medication state. However, only PD patients in the on-state quickened movement time in response to reward, while those in the off-state did not. The PD patients, in whom metabolic costs of movement tend to be higher [50], reacted more quickly to obtain reward, but only moved more quickly for the same reward when supplemented with additional dopamine. However, others have found that modulation of vigor with reward maintained in both healthy older adults and individuals with Parkinsons [51]. Movement times were measured as the average inter-key-press interval, and were faster in both groups to greater expected reward. Here, we focus on movement peak velocity, a metric we have confirmed correlated with greater energetic expenditure. Thus, it is possible that shorter inter-key-press intervals do not exact the same increase in energetic cost that would bias older adults to avoid faster movement times.

### Reducing reaction time carries a cost of accuracy

While reducing reaction time, rather than increasing movement speed, emerges as an optimal strategy to respond to increased reward in high effort environments, it nevertheless carries a risk. In our model of utility (Eq. 8), we included a speed-accuracy tradeoff on reaction time (Eq. 6). Previous work has suggested that the time before a movement can be separated into two independent phases: a preparation and an initiation phase [34,35]. While movements can be prepared rapidly, likely in the primary and premotor cortices, there is often a delay in their initiation. In the rewarded reaching task, if individuals were to initiate movements before they were adequately prepared, the odds of accurate target selection are near chance (Fig. 2A); but if individuals delay movement initiation, the probability of moving to the correct quadrant rapidly increases towards certainty [34].

Reaction times were slower overall in older adults and in the high effort environment for young adults. If we take the reaction time measured here as a sum of the preparation and initiation times, then reaction times could be lengthened by slowing either or both processes. Aging accompanies a degeneration of the nigrostriatal dopamine system, and accordingly reduced striatal dopamine transporter availability has been correlated with the slowing of reaction time in older adults [52].

Additionally, older adults see decrements in neuromuscular properties of muscle, synaptic integrity, and number of motor units, which have been linked to reaction time slowness [53]. Thus, in older adults, evidence points to slowing in both higher-level preparation and peripheral initiation contributing to their slowed reaction times.

Younger adults, when faced with increased mass and subsequent higher effort movements, may also experience slower reaction times because of changes in preparation and initiation. With higher force movements, Nagasaki et al. [47] found that both “premotor” (i.e., preparation) and “motor” (i.e., initiation) reaction times both increased, suggesting that higher forces demand increased central processing time for movement organization alongside increased time for developing the appropriate muscle tension to begin moving.

Lastly, the possibility remains that movements could have been prepared with the same rate in this experiment, but a delay in initiation reflected a more risk-averse strategy. If the effort required is higher, individuals may be increasing the delay between preparation and initiation to ensure that the movement will be successfully executed toward the correct target, avoiding unnecessarily wasted energy [34,54].

### Learning of stimulus value

Healthy aging coincides with a decreased ability to learn the value of a stimulus from its history of reward [17]. This raises the possibility that in the older group, their reluctance to increase speed of reaching may have been due to a reduced ability to learn the value of the stimulus. However, we found that in response to the rewarding stimulus, older participants decreased their reaction time by amounts comparable to the young. This suggests that lack of reward-dependent modulation in reach speed was not because of a deficit in learning value of the stimuli.

### Older and young adults executed movements similarly towards non-rewarded quadrants

In exp. 2, while older adults on average made slower reaches, we found no significant differences between the two groups when selecting peak speed in the absence of reward. These findings go against previously reported observations showing an age-dependent decrease in execution across a range of representative movements [2–8,23,55,56]. When making pointing movements, individuals adjust the speed of their movements according to the size and amplitude of the endpoint [57]. Ketcham et al. [4] reported that when reaching towards targets of decreasing size, older adults were slower and less accurate than young and were less willing to adjust the speed of their movements in response to changing task difficulty. Van Halewyck et al. [55] had young and older adults make wrist flexion movements according to different instructions and with changing feedback. In one condition, they were given visual feedback of a cursor and asked to move that cursor as quickly as possible to the center of a target. Under these constraints, older adults made slower and less continuous movements when compared to the young adult group. In a second condition, the researchers removed visual feedback of the cursor and instructed the participants to move the invisible cursor as fast as possible through the target. In this second condition, they found that older adults were able to make movements that were equally as fast and with similar variability as the young adults, suggesting that the decreased speed in reaching exhibited by older adults when given full visual feedback was not due to an inability to reach faster, but was rather a result of a change in movement strategies aimed at minimizing accuracy costs.

To best capture the relationship between effort and reward, it was vital that we minimized the cost of accuracy. The quadrants used in our experiments were of a very large size that allowed for a minimal influence of accuracy constraints. We also attempted to minimize error by removing visual feedback of the cursor during the outward portion of the movement. As long as the movement was directed towards the correct quadrant, no amount of naturally occurring signal dependent or independent noise would cause a trial to fail. These two combined factors allowed us to mitigate the cost of accuracy and instead isolate how effort and reward interact to establish vigor in older adults. They may have also mitigated the magnitude of age-dependent effects on movement speed.

### Limitations

The reward used in this study was binary. Either a reach resulted in delivery of the full reward or no reward at all. Because of this binary design, we are unable to comment on whether an effect of reward was present, but just too small to detect, on reach speed of older adults.

A few studies suggest that animals change behavior not just because of a change in reward quality, but also a change in reward rate [58]. This means that the reward landscape can be additionally manipulated by changing the frequency of reward, either by changing the relative number of rewards, or the amount of time elapsed between trials. A similar constraint implemented with older adults could further explain how they consider changing reward when establishing movement vigor.

Furthermore, we did not determine the exact energetic costs of moving faster with more effort. Changes in movement vigor were quantified in terms of absolute differences in movement duration and reaction time, in which we saw reductions in both when responding to reward (Figs 2-4). Focusing specifically on movement duration, we did not consider the exact energetic cost (in Joules) of moving *x* cm/s more quickly. That is, it remains possible that individuals were willing to allocate the same additional *n* Joules to gain reward in both the low and high effort environments; however, movement speeds simply increased less in the high effort condition because those additional *n* Joules had less of an absolute effect.

## Conclusion

We found that the metabolic cost of reaching as a function of duration and distance was elevated in older adults, and that the speed that maximized utility of reaching, defined as reward acquired minus energy expended divided by time, was slower for older adults when compared to young. When exposed to added reward, both young and older adults responded by decreasing reaction time. However, when executing the movement towards these rewards, only young adults increased their speed. When explicitly forced to reach with higher effort, a new cohort of young adults responded to reward like the elderly: in a high effort environment, the proportion of time-savings due to changing movement speed decreased while reaction time’s proportion increased. Thus, the increased metabolic cost of reaching in older adults appears to be a significant contributor to age-related movement slowing.

## Methods

### Experimental model and subject details

A total of 88 healthy subjects participated in our studies. For Exp. 1, we recruited twelve young adults (25±2 years, 6F, 6M, 66±11kg) and twelve older adults (75±8 years, 6F, 6M, 73±18kg). For Exp. 2, we recruited twenty young (26±4 years, 10F, 10M) and twenty older adults (72±6 years, 10F, 10M). For Exp. 3, we recruited twenty young adults (23±4 years, 10F, 10M). Participants were naïve to the experiments and gave written informed consent approved by the University of Colorado Boulder Institutional Review Board before participating in this protocol. All participants reported being primarily right-handed [59] and reported no issues regarding their physical or mental health. Additionally, all older adults were deemed fully mobile as evident in earning the maximum score when performing a short physical performance battery [60]. Young adult data for experiment 2 has been reported previously in Summerside et al. [28].

### Method Details

#### Experimental Tasks and Protocols

##### Experiment 1

The primary goal of experiment 1 was to quantify the effect of reaching speed on the metabolic cost of reaching. Participants sat in a chair designed to limit trunk movement and grasped the handle of a robotic arm using their right hand (Interactive Motion Technologies Shoulder Elbow Robot). The robotic handle operated similarly to a computer mouse, where movements along the horizontal plane controlled the position of a virtual cursor projected on a vertically positioned LCD monitor located at eye level (Fig. 1A). Experiments began with measurement of the metabolic rate of energy expenditure while the subjects held the handle of the robot and maintained stillness at the home position of the cursor (baseline metabolic rate). After 5 minutes of stillness, they made alternating out and back movements along the anterior-posterior (y) axis. For odd numbered trials, the movement was away from the body and on even numbered trials, the movement was back towards the body. A trial began by placing the cursor over a home circle. After a 150ms delay, a circular target (diameter=1.6cm) appeared and the participant was instructed to move the cursor to the newly projected target and stop. Participants executed their movements in this experiment according to two prescribed distances, each with five prescribed durations. The distances were 10cm and 20cm. The five movement durations used at each distance are referred to here on as very slow (VS), slow (S), medium (M), fast (F), and very fast (VF). Across all distance/duration combinations, the metabolic cost of reaching was obtained for 10 conditions per participant. The number of trials for each condition depended on reach duration and distance as well as age group (Table 1). The number of trials was selected to allow for ∼6 minutes of reaching, with faster duration conditions requiring more trials.

**Table 1.** Constrained movement duration and trial number by age and distance.

|  |  |  |  |  |  |
| --- | --- | --- | --- | --- | --- |
| Duration (ms) / Trial # | Very Slow | Slow | Medium | Fast | Very Fast |
| Young @10cm | 1000/180 | 775/200 | 500/230 | 350/260 | 125/300 |
| Older @10cm | 1100/180 | 800/210 | 600/230 | 500/245 | 250/260 |
| Young @20cm | 2050/140 | 1150/160 | 800/200 | 500/230 | 250/260 |
| Older @20cm | 2150/130 | 1250/170 | 850/210 | 550/230 | 250/250 |

Participants learned the desired duration for each condition based on two different feedback sources. The first was a training bar that would accompany the cursor along the left side of the movement path for the first four of every twenty trials. Upon movement initiation, this training bar would follow a minimum jerk trajectory towards the target indicating the prescribed reaching speed. The second source of feedback was a change of target color once the cursor made contact with the target. If the cursor arrived within 50ms of the desired duration, the target would flash yellow and deliver a pleasing tone. If the movement was too fast, the target would turn green and if the movement was too slow, it would turn gray.

Each visit began with measurement of baseline metabolic rate (holding the handle still), then five blocks of reaching at a single distance (Fig. 1B), and then concluded with another measurement of baseline metabolic rate. Each reaching block consisted of 20 practice trials accompanied with the training bar, a short 1-minute break, then an additional ∼6 minutes of reaching while wearing the nose clip and mouthpiece. Five-minute mandatory rest periods were included between blocks of reaching to allow an individual’s metabolic rate to return to rest before the start of a new block. The constrained movement duration was consistent within each block and the block order was randomized for each participant.

##### Experiment 2

The purpose of experiment 2 was to measure the effect of reward on reaching vigor. Participants were seated in a position identical to experiment 1 (Fig. 1A). A trial began with a small green circular home target (diameter=0.9cm) appearing in the center of the screen. Participants moved the cursor (diameter=0.6cm) to overlap with the home target. After overlapping for a brief 150ms, the home circle vanished, a quick audio stimulus was delivered (50ms @110Hz followed by 50ms @ 220Hz), and a larger red outer circle appeared (diameter=14cm) with its center the same as the home target. The outer ring included a small indicator located at one of four alternating locations (45°, 135°, 225° or 315° from right horizontal). The goal of the task was to move the cursor through the outer ring while staying within the quadrant containing the indicator (Fig. 2A). Once passing the outer ring, the outer ring changed color from red to gray, signaling to the participant that they should return the cursor to the center. When the cursor returned within 9cm of the center, the home target was re-illuminated to allow the next trial could begin.

In a minority of the trials (25%) a quadrant would be paired with a reward. The only requirement for receiving the reward was that the cursor crossed the 100-degree region centered on the quadrant indicator. This large region was intended to remove any differences in reach kinematics related to movement variability. The qualities of the reward stimulus consisted of a pleasing sound (50ms @880hz followed by 50ms @ 3520hz) and a visual animation of the outer ring (ring flashed yellow for 50ms and disappeared), both simultaneously delivered when the cursor crossed the outer ring. At the end of a rewarded trial, participants received 4 points, with the total accumulated points displayed on the upper right corner of the monitor.

Importantly, participants were instructed only to reach in the direction of the indicated quadrant and were informed that nothing they did beyond that would change the quantity or quality of the reward. As long as they completed the trial in the indicated quadrant and that trial was rewarded, they would receive the full reward. If participants inquired about whether they needed to perform under any time or kinematic constraints, they were told that there was no wrong way to perform the movement and to simply reach in a manner that felt natural for them. The participants were unaware of the number of trials they would be completing, only to expect the experiment to last one hour. Each participant was compensated $15 for their time with this amount being independent of any aspect of their performance in the task.

Experiment 2 began with a familiarization protocol consisting of a single block of 40 trials (10 trials to each quadrant). During these trials, participants were able to familiarize themselves with the task as well as adjust the position of the chair to ensure all four quadrants could be comfortably accessed.

The experimental protocol came after familiarization and consisted of a baseline block of 40 trials (10 trials/quadrant) followed by four experimental blocks of 100 trials (25 trials/quadrant). At the start of the experimental protocol, the participants were informed that they would no longer be receiving visual feedback of their cursor when reaching. They were also told that quadrants would now be occasionally rewarded and that as long as they reached in the indicated quadrant, they would receive the full reward.

At the start of each trial, visual feedback of the cursor was removed. The cursor re-appeared during the return movement once arriving within 9cm of the home target. There was no reward tied to any quadrant during the baseline block. In each of the experimental blocks, a single quadrant was consistently rewarded, with the location of the reward changing at the beginning of each new block.

The presentation of quadrants was completely random within each experimental block. This means that there was at any time a 25% probability that the next trial would be in the rewarded quadrant. No participants were ever explicitly informed about when a new block began, the location of future rewards, or how rewards were distributed within or across blocks.

##### Experiment 3

The purpose of experiment 3 was to directly assess the effect that added effort has on reward responsiveness on reaching vigor. The task was almost identical to the task from experiment 2 outside of a few changes: *(i)* participants completed 80 trials in the baseline blocks before completing the four blocks of 100 trials (25 trials/quadrant); *(ii)* the distance from the central home circle to the edge of the target ring was also reduced to from 14cm to 10cm; and *(iii)* participants had visual feedback of the cursor throughout the trial, though were instructed that the only criterion for success was to cross the cursor anywhere along the 100° quadrant.

To gauge how the response to rewarded quadrants changes with added effort, participants performed the above experimental protocol twice – once with low effort (0 kg added mass), and once with high effort (3.63kg; 8lbs), physical mass added to the robotic arm. The order of the effort conditions was randomized and counterbalanced across participants. For 6 participants, the completion of the task with low and high effort occurred on two separate days; the remaining 14 participants performed both conditions on the same day with at least 10min of rest between each condition. Between the low and high effort conditions, each participant experienced a total of 160 baseline trials, 200 rewarded trials, and 600 nonrewarded trials.

#### Measuring Metabolic Cost

In experiment 1, we quantified effort by measuring metabolic cost as a function of movement distance and duration. Metabolic cost was measured via expired gas analysis (ParvoMedics, TrueOne2400).

Participants wore a nose clip and breathed in and out of a mouthpiece throughout all reaching bouts in experiment 1. This allowed us to measure how the rates of oxygen consumption and carbon dioxide production changed across conditions. To minimize the thermic effect of food on metabolic rate, all sessions were conducted in the morning with participants arriving having fasted overnight. The metabolic cart was calibrated at the start of each visit according to certified gas mixtures as well as a range of flow rates from a 3-litre calibration syringe. Baseline resting metabolic rate was measured while participants sat quietly in the chair holding the robotic handle. Baseline resting trials were taken at the start and end of the visit.

We calculated gross metabolic rate (*ė*_*gross*_) and resting rate (*ė*_*o*_) in watts (J/s) using the rate of oxygen consumption and carbon dioxide production according to the Brockway equation [61]. We only included conditions where the respiratory exchange ratio was between 0.7 and 1.0, indicating aerobic respiration. We calculated net metabolic rate during each condition by measuring the gross metabolic rate (*ė*_*gross*_) averaged over the last three minutes of reaching in each condition and subtracting the lower of the two metabolic rates measured during seated rest (*ė*_*o*_) at the beginning and end of the visit.

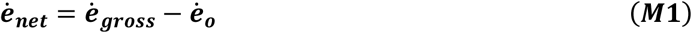

Our goal was to calculate the metabolic cost of moving (*ė*_*move*_). The net rate in Eq. 4 represents the combined cost of moving (*ė*_*move*_) and the cost of not moving between each trial (*ė*_*ITI*_) above rest:

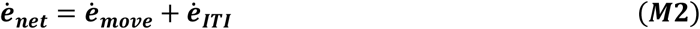

If we assume that the cost of waiting between trials is equal to the cost of rest, then *ė*_*ITI*_ is equal to zero. We can then localize the metabolic cost of moving during each trial according to the equation:

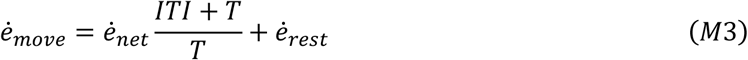

Here *ITI* represents the average length of the inter-trial-interval (ms) and *T* represents average movement duration (ms). We then fit the calculated metabolic rate of moving according to Eq. 1.

To calculate the total energy required of each movement, we multiplied the metabolic rate of moving by the movement duration and refer to this calculation as the metabolic cost of reaching (CoR).

### Quantification and Statistical Analysis

#### Data analysis

Mediolateral (x) and anteroposterior (y) positions of the handle were recorded at 200Hz. With these signals we calculated radial position, which was then smoothed (Fourth order Butterworth filter with cutoff frequency of 10Hz.). Instantaneous speed and acceleration were obtained through differentiating smoothed radial position. Jerk was calculated independently along the x and y axes by triple differentiating the position signals along those axes.

Reaction time was calculated as the difference in time between when the audiovisual start stimulus was delivered and movement onset. Movement onset was identified using a threshold based on both radial speed of 0.05 m/s and radial acceleration of 0.05 m/s^2^. In experiment 1, movement offset was determined as the last moment that the speed exceeded a threshold of 2.5cm/s. In experiment 2, offset was determined as the moment of maximal radial position from the home. Peak instantaneous outward speed was identified between movement onset and movement offset.

Movement duration was calculated as the difference in time between movement onset and offset. Total distance was measured as the difference in position between movement onset and movement offset. Inter-trial-interval (ITI) was measured between each trial as the time between movement offset of the current trial and movement onset of the subsequent trial. This meant that ITI represented the combined time spent repositioning the cursor for the next trial and the reaction time of that same trial.

Trials in experiment 2 with reaction times greater than 700ms or with crossing-point distances outside of the 100° quadrant were removed from analysis. Across all young participants this accounted for an exclusion of 0.46% of trials (43 for reaction time and 2 for crossing the ring in the incorrect quadrant). For the older participants, a total of 4.32% of trials were excluded (332 trials for reaction time and 14 trials for crossing the outer ring in the incorrect quadrant).

Trials in experiment 3 were removed if reaction times were >700ms or <100ms. Total, this excluded 0.98% of all trials (165/16,800). Trials would have been excluded if the cursor did not cross the ring within the 100° arc; however, no trials were unsuccessful in this manner.

#### Statistical Analysis

##### Experiment 1

All young adults were able to complete each of the five durations at each of the two distances. This resulted in a total of 120 metabolic measurements for the young adults. Two older adults were unable to complete a single condition resulting in a total of 118 metabolic measurements for the older adult group. To determine the effect of age on the metabolic rate of moving, we implemented a linear mixed effects regression predicting the log transform of *ė*_*move*_ as a function of a binary age indicator (older=1), a binary distance indicator (10cm=1), the log transform of average velocity (continuous, seconds) and the resting rate of each participant (continuous, Watts). We performed another linear mixed effects regression to predict movement endpoints in x- and y-axes and sum of jerk squared 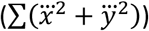 and sum of squared joint torques as a function of a binary age indicator (older=1), a binary distance indicator (10cm=1), and average velocity (continuous, seconds). For the sum of jerk squared measure we used the log transform of velocity and sum of jerk squared. Differences in resting rate between the young and older adults were explored using an independent t-test with each individual’s resting rates averaged across both visits.

For each age group, we described the metabolic landscape of reaching as a function of distance and duration according to equation 1. All parameter values were obtained via bootstrapping (*nls* function and *boot* package available in *R* version 3.3.3). The young and older adult datasets were sampled over 10000 replications with each replication consisting of 120 points sampled with replacement (12 participants per age group, 10 conditions per participant).

We estimated the mass of the upper arm, lower arm, and hand using previously published estimates according to age, sex, and body mass [25,26]. Differences in segment mass between the young and old groups were compared using two-sample independent t-tests.

For each age group and movement duration, we used the mean and standard deviation of subject endpoint error to calculate the probability that the endpoint was within the target radius (0.8cm). Probability of reward was then described as a function of reach duration according to equation 5.

Reaction time probability of reward was described with a similar equation (6). Without data to fit, the parameters *c*_0_ and *c*_l_ were determined assuming a non-zero probability of reward when reaction time was zero, and a 100% probability of reward at reactions times of approximately 400ms. Theory predictions remained conceptually the same across a range of parameter values.

##### Experiment 2

The effect of reward for every individual was quantified by comparing the average reaction time and peak speed of each reaching movement towards a quadrant when it was rewarded minus when that same quadrant was not rewarded. Reaction time and peak speed were modeled using a linear mixed effects regression with age, reward, and trial number as predictors. Age was binary, trial number was continuous, and reward status was binary. Additional predictors included a reward-by-age, age-by-trial, and reward-by-trial interactions. Rewarded movements were further compared to non-rewarded movements in the trials immediately before and after for both age groups using paired t-tests. Response magnitudes in all trials before and after reward were compared between age groups using a two-sample independent t-test.

##### Experiment 3

The primary kinematic measures of interest were reaction time, peak outward velocity, movement duration, and movement vigor. To describe the both the individual and interacting effects of effort and reward, we constructed linear mixed effects regression models predicting each of the kinematic outcomes (peak velocity, reaction time, movement duration, and vigor). Fixed effects in each model were a binary reward predictor (rewarded = 1), a binary effort predictor (high effort = 1), and a reward-effort interaction. Between-subject variation was incorporated as a random intercept. All valid reaches were included when fitting the model. A statistical threshold of α = 0.05 was used for all comparisons. All p-values were adjusted for multiple comparisons using the Holm-Bonferroni method on a family-wise basis.

Results from the above statistical analyses are reported in the main text of the results section as well as in the supplementary results. All statistical thresholds were conducted at a significance level of α=0.05. Descriptive statistics are reported as mean ± standard error.

Computational model of movement time and reaction time: To investigate this idea that increases in movement effort should predispose individuals to emphasize reacting faster as opposed to moving faster, we ran simulations using the utility of movement as a cost function:

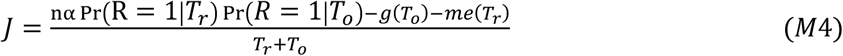

We iterated over a series of reward values (*α* = {20,21,22 …,200}), reward scaling coefficients (*n* = {0.8, 0.9, 1.0, 1.1, 1.2}), and effort scaling coefficients (*m* = {0.8, 0.9, 1.0, 1.1, 1.2}) to perform a constrained search (*fmincon* function in MATLAB) for the reaction times (*T*_*o*_) and movement times (*T*_*r*_) that maximized utility (*J*) (Eq. 1) for a given reward. Fitted parameters in the metabolic effort (Eqs. 2 & 4) and reward probability (Eq. 5) functions were set to those of the young and older adults as applicable.

### Data and software availability

All data used to draw the results presented in this report, including the R code necessary to analyze these data can be freely accessed at the following location: https://osf.io/cfm9v/?view_only=df7ccb37d27a4aeab01c1e2f83c40e98

