## Supplemental Figure 1 for "Effort cost of reaching prompts vigor reduction in older adults"

### Supplemental Results

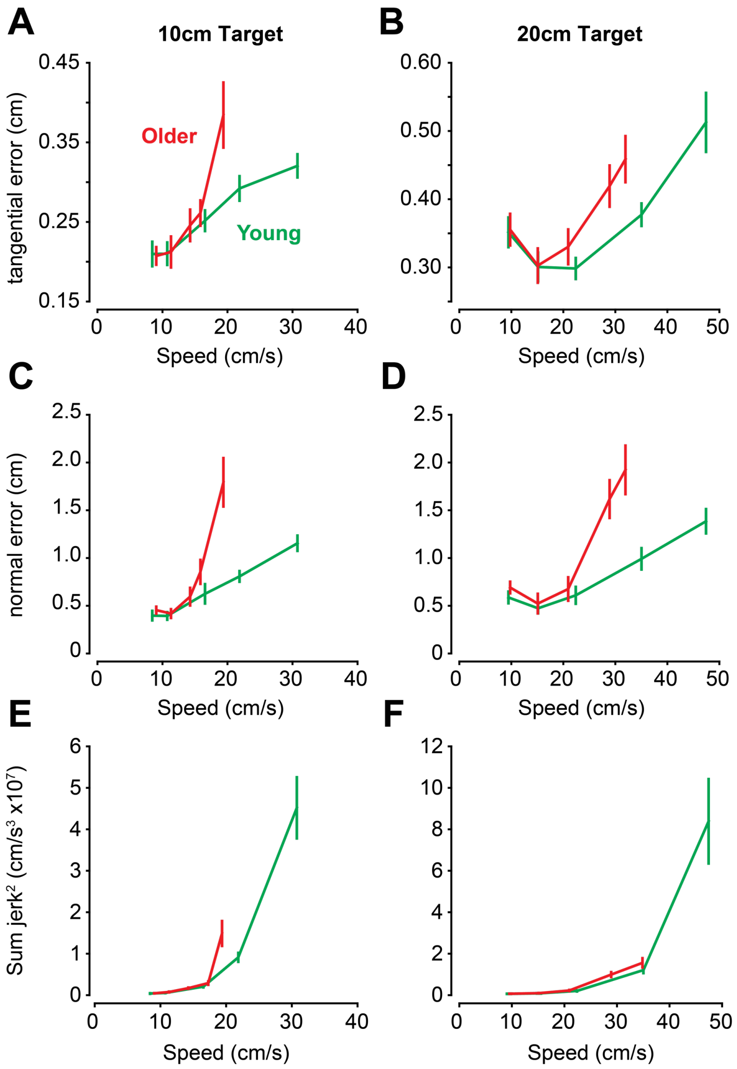


**Figure S1.** Effect of increasing speed on mean endpoint error and sum of jerk$^2$ in young (green) and older (red) adults. A,B) Young and older adults similarly increased endpoint error along the tangential-axis as a result of increasing speed. C,D) Increasing speed also led to greater endpoint error along the normal axis. This effect of speed on endpoint error along the normal axis was greater in older adults. E,F) Effect of increasing speed on the sum of jerk$^2$. This speed effect was indistinguishable between age groups. Bars represent ±SEM.
